## Supplementary material for "CABaNe, an automated, high content ImageJ macro for cell and neurite analysis": Figure 2-1 User guide

First, images have to be sorted in conditions files. This can be either done automatically, by running CABaNe, setting parameters, and only choosing “Sort”. To use the “Sort” function, filters have first to be tested with “test_filter”, in a mock folder, to insure the filters are fitted to the images. Folder selection is done in a separate window once you click ok: the first folder is the one containing the images, and the second the one that will contain **the parent file** for sorted images and their analysis. This will creates folders for analysis, store valid images and store rejected images. Images can be automatically sorted in condition if their name formatting is X-Y(info).tif, with X and Y are a well position. Otherwise, images can be stored in a general folder. Alternatively, the user can do it manually, by following the architecture presented in **Figure 2-3**. Only the first 3 layers of folder need to be created, up to condition A, B,C… Names of folders can be changed. Both channel of cell and nuclei (and eventually an intensity marking) for each images must be stored separately in the folders. Identify a string of characters unique to each channel.

Once images are sorted, the macro can be ran. If filtering parameter need to be tuned, it is possible to use the “test_filter” option, choosing the folder containing images, the start and end condition of testing, as well as the number of images (cell + nuclei) to test per conditions. CABaNe will run mock analysis, and present the results of masks resulting of selected parameters, as presented in **Figure 4**. It is possible to either cancel or continue viewing subsequent well. Once it reach the final well, the base interface will reappear, keeping previously selected parameters.

Once the images are sorted and parameters set, the main function of CABaNe, analysis, can be started, after selecting the parent folders for images and results (see **Figure 5**).

After clicking ok, CABaNe will start running in background. Depending on the number of images and cells, the analysis can be quite long, so it is recommended to either run little bits of the global set, run in on a separate computer, or overnight. The log window indicate overall progression, for each image. While it is running, it might be intricate to use the computer, due to the nature of ImageJ macros, as they are given priority when opening a new window.

A message will indicate end of the run. It is then possible, depending on needs, to go to the files to get the results or to check the analysis.

Results are located in the bottom “Results” folder (see **Figure 5**), containing single cell tables and summary for each field of view. Subsequent field of view summary are built upon previous summary of a same conditions, and so the summary of the last field of view is actually is a summary of the whole condition. It is possible to either manually retrieve the data, or to use the bonus python script, by modifying the code to include the path of the parent file of the experiment. Result file have to be named “Result” and conditions well names such as “A01”, B04” and so on.

To check the analysis, it is possible to check cell and nuclei segmentation in the image, opening the .tif in the “Segmentation checks” folder. For single cell pathing, images in “checks” represent the segmentation and pathing of single cells. To choose which channel to see, use the Channel tool in image J. Channel 1 is the cells contouring after watershed. Channel 2 is the nucleus segmentation. Channel 3 is the skeleton of cells. Channel 4 is the result of the masks on the cell channel. Channel 5 is the base image. Channel 6 is the body of each cell, after watershed.
