## Supplementary material for "CABaNe, an automated, high content ImageJ macro for cell and neurite analysis": Figure 2-2 Troubleshooting

Table 5: Known issue for CABaNe. Each is linked to example, theory on what is happening, and a potential solution.

| **Issue** | **What is happening** | **What is the solution** |
| --- | --- | --- |
| Error message “metada.txt path not found” | The windows error is typical of a wrong folder configuration. | Create or rename a folder in the “result” folder to match the “Images” folder. See **Figure 9** for more information, and **first paragraph of user guide**. |
| Error message “ROI manager: The list is empty” | CABaNe failed to detect any ROI for cells, bodies, or nuclei. This can also be due to an empty image. | Use the “sort” function to prevent this from happening, or a manual check. The “test_filter” option can also be used to check if on current images, the cell detection issues is caused by the image having no cell, or if the filters are wrongly set, leaving to a poor cell detection. See **second paragraph of user guide**, and **Figure 4**. **See next issue.** |
| Filtering of images is well tuned but cell detection limit is wrong, cells are ignored. | Accepted cell range is too strict, and exclude cells. | Try widening (here, lowering the minimum cell size) the cell detection parameter. This can also happen with nuclei, or bodies (that are indexed on cell size to limit the number of parameters, but it is easy to change should it be needed.). Used type of detection could also be too discriminative, in which case you should switch to a more tolerant one: Mean > Triangle > Minimum |
| Imprecise detection, different cells are merged together | Used filter is not discriminative enough | Try using a more discriminative filter if possible, on either nuclei, cell, or body, depending on which type of detection is merging. Discrimination: Mean < Triangle < Minimum. You can change the filter in the code if you have specific needs. Remember to use “test_filter” before analysis to check your filters. |
| Partial detection in dense area. | Cells are a bit too dense in the image, creating 3D structure, which impaired the focus. Therefore, some part of the cells are out of focus. | This will make it harder for CABaNe to detect the cells from the background, especially if there is noise. The user can try using less discriminative filters, using images with less cells or a better focus, or consider that the data acquired is sufficient to extrapolate. |
