## Supplementary figures and images for "CABaNe, an automated, high content ImageJ macro for cell and neurite analysis"

### Figure 2-3 folder architecture

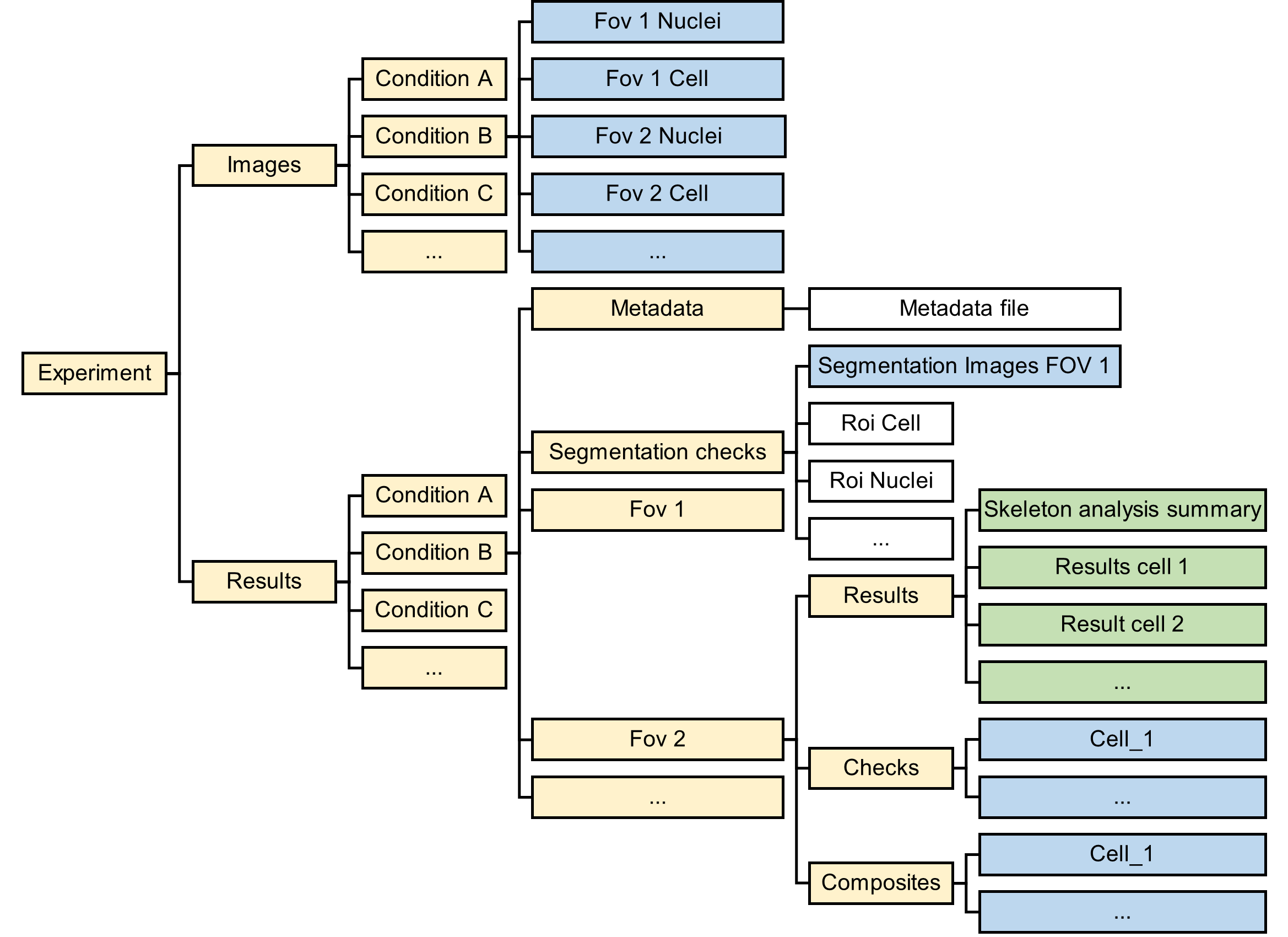

### Figure 5-1 segmentation comparison

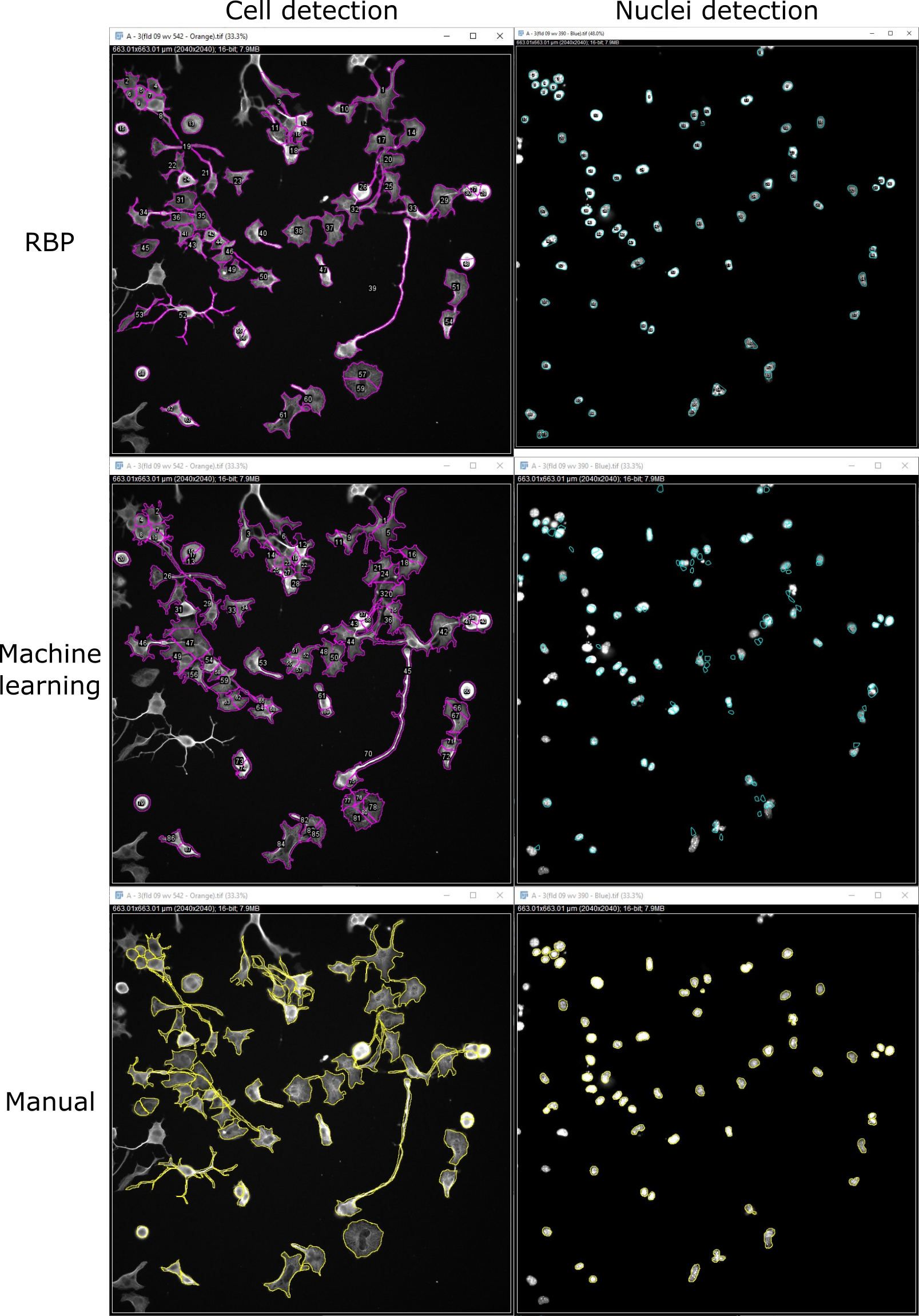
