## Supplementary material for "CABaNe, an automated, high content ImageJ macro for cell and neurite analysis": PYTHON_CODE

import os
import pandas as pd


#Input path to parent folder with your “Result” folder
Filepath= str('C:/Users/')
Startingrow=    0
Endingrow=      8                                                              #Ending is the one you wanna stop before, counting of list start at 0
Startingcolumn= 0
Endingcolum=    12
row=            ["A", "B", "C", "D", "E", "F", "G", "H"]
column =        [1, 2, 3, 4, 5, 6, 7, 8, 9, 10, 11, 12]
to_file= "".join([Filepath, '/SUMMARY_RESULTS.xlsx'])                           #join the string to create a bigger string
r=              Startingrow
c=              Startingcolumn
for r in range(Startingrow,Endingrow):                                          #looking at all rows (since data is stored in rowcolumn format)
    for c in range(Startingcolumn,Endingcolum):                                 #looking at all column
        if(column[c]<10):
            Well= str(row[r]) + "0" + str(column[c])
        else:
            Well= str(row[r]) + str(column[c])                                      #to create name of the well we are looking for (name of the folder)
        Currentfile= Filepath + '/RESULT/' + Well                               #as indicated before, here we use the path, plus result (where the result are) plus name of the well
        try:
            Listeresultdossier= os.listdir(Currentfile)                             #'read' the file of before, to get access to all the elements
            PositionResultfile= len(Listeresultdossier)-3                           #find the correct file to implement
            from_file= Currentfile +'/'+ Listeresultdossier[PositionResultfile] +'/Results/_Skeleton_analysis_summary.csv' #engineering of the complete name of the path by adding the typical path to result
            data=   pd.read_csv(from_file,delimiter=",")                            #understand as 'read the csv using this delimiter'
            with    pd.ExcelWriter(to_file,mode= 'a', if_sheet_exists='overlay') as writer:     #overlay : set to replace if there is anything on the existent sheet
                pd.DataFrame(data).to_excel(writer, header=False, index=False, sheet_name= Well, startrow= 2, startcol=1) #header and index are not data of value, so they are not copied into the sheet. Startrow/col define emplacement of copying in the file.
                print(Well)
        except:
            pass
            print(Well+' do not exist')
SystemExit()
